# Optical genome mapping identifies source-associated structural variant differences across early-passage human iPSCs

**DOI:** 10.64898/2026.05.29.728843

**Authors:** Leila Namvar, Kamilla Sedov, Madison James Yang, Rebecca Hermosillo, Faria Zafar, Birgitt Schüle

## Abstract

**Background:** Induced pluripotent stem cells (iPSCs) are an important model for studying human diseases *in vitro*. However, previous studies have shown that iPSC reprogramming and extended cell culture can introduce genomic structural variants (SVs). Technologies like karyotyping, CNV microarrays, and whole-genome sequencing have limitations in resolution, sensitivity, or the ability to detect large and complex structural variants compared to optical genome mapping (OGM). OGM is a genome-wide structural variant detection method that analyzes fluorescently labeled ultra-high-molecular-weight DNA molecules to identify copy-number and balanced rearrangements. At sufficient coverage, OGM can detect SVs at approximately ≥2 kbp and identify mosaic events supported by molecule-level evidence, offering higher resolution than conventional karyotyping or SNP-array-based QC.

Here, we compared iPSC clones derived from peripheral blood mononuclear cells (PBMCs) and fibroblasts (FBCs) to determine whether starting somatic cell source is associated with differences in structural variant burden and SV-type profiles after nuclear reprogramming into iPSCs.

**Results:** We analyzed 73 low-passage iPSC clones generated from 25 parental lines using OGM. Compared with PBMC-iPSCs, FBC-iPSCs showed higher SV burden with the enrichment of duplications ≥100 kbp, more frequent overlap with protein-coding genes, fragile sites, and recurrent chromosomal hotspot regions. In contrast, PBMC-iPSCs showed fewer SVs overall, and a higher proportion of clones without detectable clone-specific SVs.

**Conclusions:** OGM provides a high-resolution approach for post-reprogramming genomic quality control by detecting clone-specific structural variants at approximately ≥2 kbp, including events below the resolution of conventional cytogenetic and SNP-array-based assays. In these early passage iPSCs, SVs overlapped protein-coding genes, fragile sites, and recurrent culture-associated chromosomal regions, underscoring the need for clone-level genomic assessment before downstream applications. FBC-derived iPSCs showed a higher SV burden, including more frequent and larger duplications, whereas PBMC-derived iPSCs more often lacked detectable clone-specific SVs. These findings suggest that PBMC-iPSCs and FBC-iPSCs can differ in post-reprogramming SV profiles and support the use of OGM as a QC strategy during iPSC generation and selection.

## BACKGROUND

Human induced pluripotent stem cells (iPSCs) are reprogrammed somatic cells, commonly from blood or skin, that can differentiate into all three germ layers[1]. iPSCs hold great potential for both research and therapeutic applications, serving as models for human diseases. Reprogramming somatic cells to iPSCs requires the expression of key Yamanaka transcription factors.

Early reprogramming methods, such as retrovirus or lentivirus, rely on viral vector integration but often result in low reprogramming efficiency, accumulation of point mutations, and chromosomal aberrations[2]. In contrast, non-integrating approaches, such as episomal and mRNA-based delivery systems, reduce the risk of permanent vector integration and are therefore preferred for generating iPSC lines intended for disease modeling or downstream translational applications[3][4]. However, even non-integrating reprogramming can be associated with de novo or selected genomic changes during reprogramming and early culture. This distinction is important because paired analysis of parental and iPSC genomes can help distinguish clone-specific structural variants that emerge or become enriched during reprogramming from variants introduced by the delivery system or already present in the starting somatic population.

Although iPSCs are expected to retain the genome of the starting somatic cell population, including preexisting genetic variants or mosaic abnormalities, additional genomic changes can arise or become enriched during reprogramming and subsequent culture[5]. These changes range from single-nucleotide variants and small insertions or deletions to larger copy-number changes and structural variants (SVs). SVs can occur across the genome, but certain regions, including common fragile sites, are particularly susceptible to replication stress, DNA breakage, and rearrangement[6].

Standard approaches for detecting structural variants (SVs) differ in resolution, genome coverage, variant classes detected, and suitability for routine quality control. Conventional cytogenetic methods can identify large chromosomal abnormalities but have limited resolution. Karyotyping detects aneuploidies and large balanced or unbalanced rearrangements, typically at megabase-scale resolution, whereas FISH provides higher, locus-specific resolution but requires predefined probes and does not offer unbiased genome-wide detection[7][8]. Array-based methods, including copy-number variation (CNV) microarrays and SNP arrays, provide genome-wide detection of copy-number gains and losses, but have limited breakpoint precision, do not reliably detect balanced rearrangements, and have reduced sensitivity for low-level mosaicism[7][9]. Short-read whole-genome sequencing can detect single-nucleotide variants, small insertions and deletions, copy-number changes, and some SVs, but detection of large, repetitive, balanced, or complex rearrangements remains challenging because short reads often cannot be uniquely mapped across repetitive or structurally complex regions[10]. Long-read sequencing addresses many of these limitations by generating reads that can span repetitive regions and complex breakpoints, enabling more complete SV discovery and improved breakpoint resolution. However, long-read sequencing remains more costly, lower-throughput, and analytically more complex than routine cytogenetic or array-based QC approaches, which can limit its use for large-scale iPSC clone screening[11].

Optical genome mapping (OGM) provides an orthogonal, genome-wide approach for structural variant detection that complements conventional cytogenetic, array-based, and sequencing-based methods. OGM can detect SVs typically ≥2 kbp and can identify both balanced and unbalanced rearrangements, including those present at low-frequency variants, which are often challenging for other technologies to detect [12]. Because OGM can detect large and complex rearrangements, balanced events, and mosaic variants that may be missed by karyotyping, FISH, or SNP-array-based QC, it provides a high-resolution cytogenomic approach for post-reprogramming assessment of iPSC genomic integrity.

In this study, we asked whether non-integrating reprogramming introduces SVs in early passage iPSCs and whether SV patterns differ by starting somatic cell type. We analyzed 73 low-passage clonal iPSC lines generated from 25 donors, 13 peripheral blood mononuclear cell (PBMC) parental lines, and 12 fibroblast cell (FBC) parental lines. Each clone was compared to its matched parental cell line to assess SV burden and SV-type profiles, and to evaluate overlap with genes, fragile sites, and recurrent chromosomal regions.

## METHODS

### PBMCs and Skin Fibroblasts

PBMCs and FBCs were received directly from the Stanford Alzheimer’s Disease Research Center (ADRC) stem cell repository or the Michael J. Fox Foundation (MJFF). PBMCs were collected in a Ficoll density gradient separation tube. Fibroblasts were generated using the skin punch biopsy explant culture protocol [13] or enzymatic treatment to derive primary human fibroblasts.

Reprogramming was performed internally at Stanford, by AlStem Services, or Pathways to Stem Cell Science (Table S1). The parental cell lines were reprogrammed into iPSCs from FBCs or PBMCs using non-integrating episomal plasmid- or mRNA-based delivery of reprogramming factors encoding OCT4, SOX2, KLF4, and L-MYC. Emerging colonies with iPSC-like morphology were selected and expanded as individual clones. Clone quality control included pluripotency marker assessment, mycoplasma testing, and sterility testing, with reports confirming the absence of bacterial, fungal, and mycoplasma contamination.

Pluripotency for all clones was confirmed using characterization assays, including immunostaining for TRA-1-60 and OCT4, and the assessment of alkaline phosphatase (AP) activity (**Figure S1 and Figure S2**). These assays were performed following initial clone picking to confirm pluripotency. Up to three iPSC clones per donor were thawed, maintained on Matrigel-coated Falcon 6-well plates (354277, Corning; 08-772-1B, Fisher Scientific) in mTeSR+ (100-0276, STEMCELL Technologies), and transitioned to StemFlex medium (A3349401, Thermo Fisher Scientific) for expansion. Routine passaging was performed using ReLeSR (05872, STEMCELL Technologies) at a 1:15 split ratio, with cultures maintained between passages 4–10, and morphology remaining consistent throughout.

Following the *Bionano Prep Sp-G2 Frozen Cell Pellet DNA Isolation Protocol-CG-0004, Rev E*, cell pellets from up to three iPSC clones per donor were collected between passages 4 and 10 (1.2–1.5 × 10D cells per vial) for OGM. The cells were deposited at the National Centralized Repository for Alzheimer’s Disease and Related Dementias NCRAD.

### Optical Genome Mapping Sample Processing and QC Analysis

Ultra-high molecular weight (HMW) genomic DNA was extracted using the *Bionano Prep Sp-G2 Frozen Cell Pellet DNA Isolation Protocol-CG-0004, Rev E.* DNA was labeled with DLE-1 using the Bionano Prep *DLS-G2 protocol CG-30553-1*.

Samples were loaded onto Bionano flow cell chips (PN 20440) and imaged on the Saphyr instrument, targeting 1500 Gbp (∼400× coverage). Guided and dual analyses compared parental and iPSC samples using Bionano Access (v1.8.1.1 or v1.8.2) aligned to a T2T reference genome.

Sample-specific molecule support thresholds were derived from effective coverage: homozygous SVs required coverage × 0.05 and heterozygous SVs coverage × 0.025 (∼5–12 molecules). Samples were required to meet predefined QC thresholds for effective coverage, molecule N50, map rate, and label density (Table S2) before downstream analyses (Bionano Manual CG-30223).

### Data analysis

SV data were analyzed using Bionano output fields mapped to standardized columns: RefcontigID1 (chromosome), RefStartPos (SV start), RefEndPos (SV end), and Type (SV type). SV size, VAF, and gene overlap were used as provided. “Present in % of BNG control samples” was used for control database frequency, “Found in control sample molecules” for parental molecule count, and “Found in self molecules” for self molecule count.

The dataset was sorted by SV size and organized by SV and type, and by chromosome location to compare SVs across clones. SVs present in all three sister clones were removed, while SVs shared between two of the three clones were retained. SVs present in all three sister clones were removed to reduce inclusion of parental variants or recurrent mapping artifacts not detected above threshold in the parental sample, whereas variants shared by two clones were retained as potentially clone-enriched or early reprogramming-associated events and annotated separately.

### Statistical analysis

All analyses used SV calls identified by optical genome mapping after clone–parent dual analysis and study-specific QC filtering. Unless otherwise stated, tests were two-sided and performed at α=0.05. SV counts were summarized as mean, median, and interquartile range (IQR). Graphics and analyses were performed at the clone level, with sensitivity analyses accounting for donor-level clustering.

### SV burden per clone

Total SV burden per clone was defined as the sum of insertions, deletions, duplications, and translocations (SV_total = ins + del + dup + tra). Because SV_total was a non-negative, right-skewed count variable with many zeros, group differences (PBMC-iPSCs vs FBC-iPSCs) were evaluated using the two-sided Mann–Whitney U test (distribution-robust) and the two-sided Welch’s t-test (parametric, unequal variances; secondary analysis).

To estimate count-scale effect sizes while accounting for overdispersion, SV_total was modeled using negative binomial regression with cell source (PBMC vs FBC) as the primary predictor. Effect sizes were reported as incidence rate ratios (IRRs) with 95% confidence intervals (CIs). To account for multiple clones per donor, the donor was included as a cluster-robust (sandwich) variance estimator or as a donor-level random intercept in a mixed-effects negative binomial model (sensitivity analysis).

Model fit and overdispersion were assessed by comparing Poisson vs negative binomial residual deviance and dispersion diagnostics.

### SV-type composition

Differences in SV-type composition between PBMC-iPSCs and FBC-iPSCs were assessed using a 2×4 chi-square test of independence on pooled SV classes (insertions, deletions, duplications, translocations). Standardized Pearson residuals identified classes contributing most to the association (e.g., enrichment of duplications in FBC-iPSCs).

Clones with SV_total = 0 were classified as SV-free, and proportions were compared using a two-sided Fisher’s exact test due to small, expected counts.

### SV size comparisons

SV sizes (bp) were summarized by group and SV class (insertion, deletion, duplication); translocations were excluded because the breakpoint-to-breakpoint size was not defined. Sizes were binned as <10 kbp, 10–<50 kbp, 50–<100 kbp, and ≥100 kbp. Between-group comparisons of continuous sizes used Mann–Whitney U tests due to skewed distributions, and binned distributions were compared using chi-square tests when expected counts permitted.

P-values were unadjusted for prespecified primary outcomes (SV_total burden and SV-type composition); all other analyses were exploratory and interpreted accordingly.

### Overlapping genes

SV-overlapping protein-coding genes were categorized based on known disease associations, gene constraint, and relevance to neurological or developmental phenotypes. These annotations were used to prioritize clones and were not intended for clinical classification of SVs. Genes overlapping between dual analyses were evaluated using NCBI, OMIM, ClinVar, gnomAD, MGI, and MedGen. Gene biotype was determined via NCBI, and non-coding genes and pseudogenes were excluded [14][15][16][17]. OMIM provided phenotype, cytogenetic, and inheritance data, and ClinVar provided disease variant classifications; definitive disease association was not always assignable due to incomplete gene characterization.

Gene constraint was assessed using gnomAD LOEUF scores (>1 = low constraint; ≤0.6 = loss-of-function intolerance). Functional effects of homologous deletions were evaluated with MGI, and diseases were categorized as neurodegenerative, neurodevelopmental, or both using MedGen. Statistical analyses included Poisson rate comparisons for clone-normalized gene burden and Fisher’s exact tests to compare proportions of high-priority or disease-associated gene overlaps between PBMC-iPSC and FBC-iPSC (two-sided).

### Fragile sites and common genetic hotspots

Common genetic regions were identified using Baker et al. (2016) [18]. Associated diseases were identified based on either the chromosomal locations or the associated genes. Fragile sites were annotated by comparing SV chromosomal locations with those previously described [19]Genes associated with each fragile site were identified using the Human Chromosomal Fragile Sites HumCFS database [20]. All fragile site-associated genes were verified using the UCSC Genome Browser[21]. SV coordinates overlapping fragile sites were compared with associated gene locations to determine whether the distance was within 100 kbp. Genes located >100 kbp away were not considered affected. SVs overlapping fragile sites were evaluated for disease associations based on chromosomal location, implicated genes, and the fragile site to determine potential links to cancer.

## RESULTS

### PBMC and FBC reprogramming and iPSC clone generation

PBMC and FBC samples were obtained from 25 donors (ages 30–76 years) (**Table S1**), including individuals with AD, PD, ataxias, and healthy controls. For each donor, the parental somatic cell line was reprogrammed into up to three clonal iPSC lines using non-integrating episomal or mRNA-based methods, yielding a total of 73 clones (39 PBMC-iPSCs and 34 FBC-iPSCs).

### Paired OGM workflow for clone-parent SV comparison

We built a customized OGM pipeline to evaluate genomic stability in low-passage iPSC clones (**Figures 1 and 2**). Parental cells and up to three matched iPSC clones per donor were profiled to generate molecule maps and SV calls. Each sample was first mapped to the T2T reference genome to produce a Guided Assembly genome map. Each iPSC clonal assembly was compared with the parent assembly via Dual Guided Assembly (parent = control; clone = case), creating a list of clone-specific SVs (**Tables S4 and S5**).

**Figure 1:**
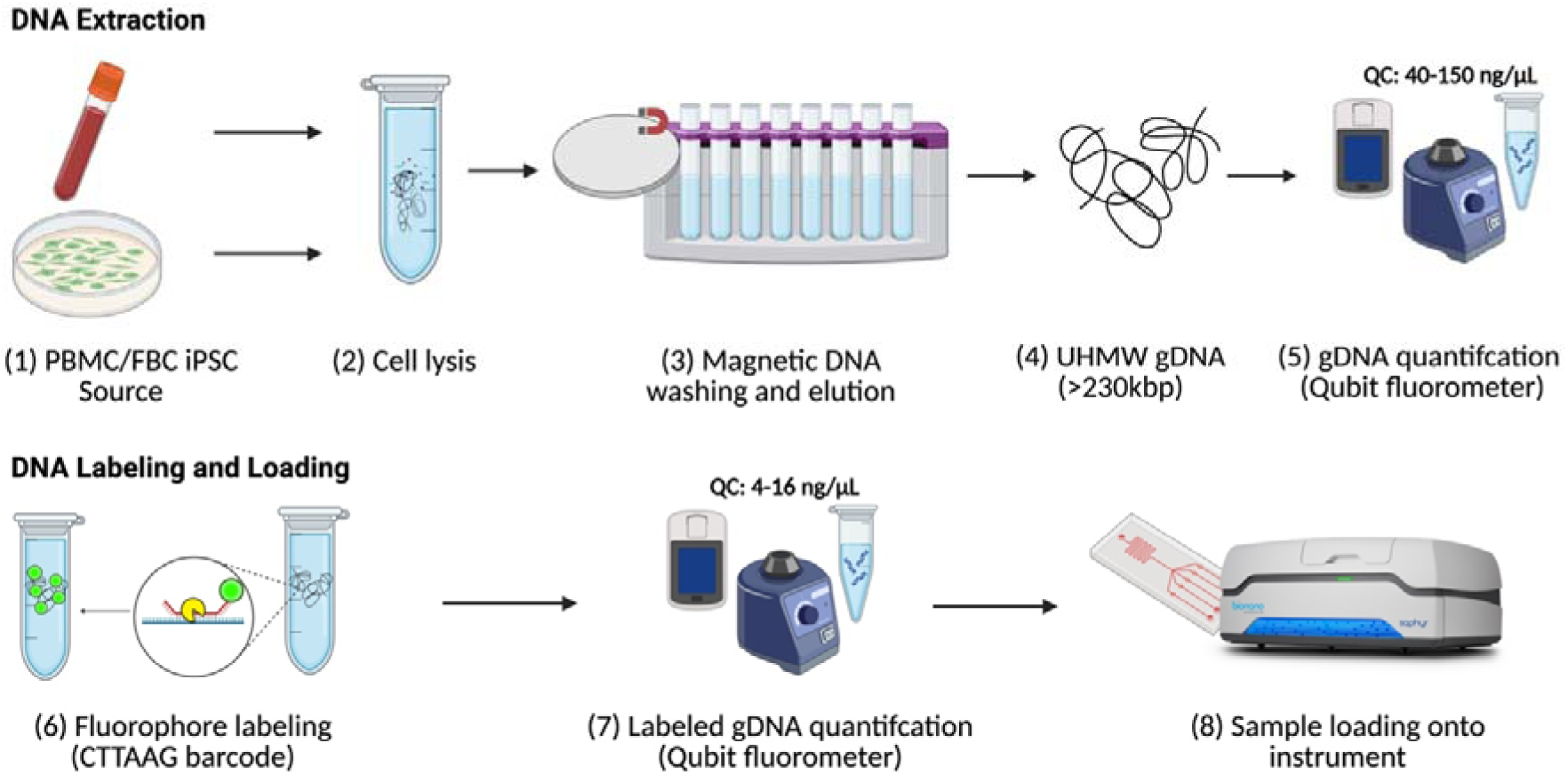
Overview of Extraction and Labeling Steps in Sample Processing. gDNA was extracted from lysed cells, washed, quantified by Qubit, labeled, validated, and then run on the Bionano Saphyr chip to generate reads (n= 73 iPSC clones, 39 PBMCs, and 34 FBCs). DNA extraction and labeling used the *Bionano Protocol-CG-0004, Rev E, and DLS-G2 Protocol* CG-30553-1.

**Figure 2:**
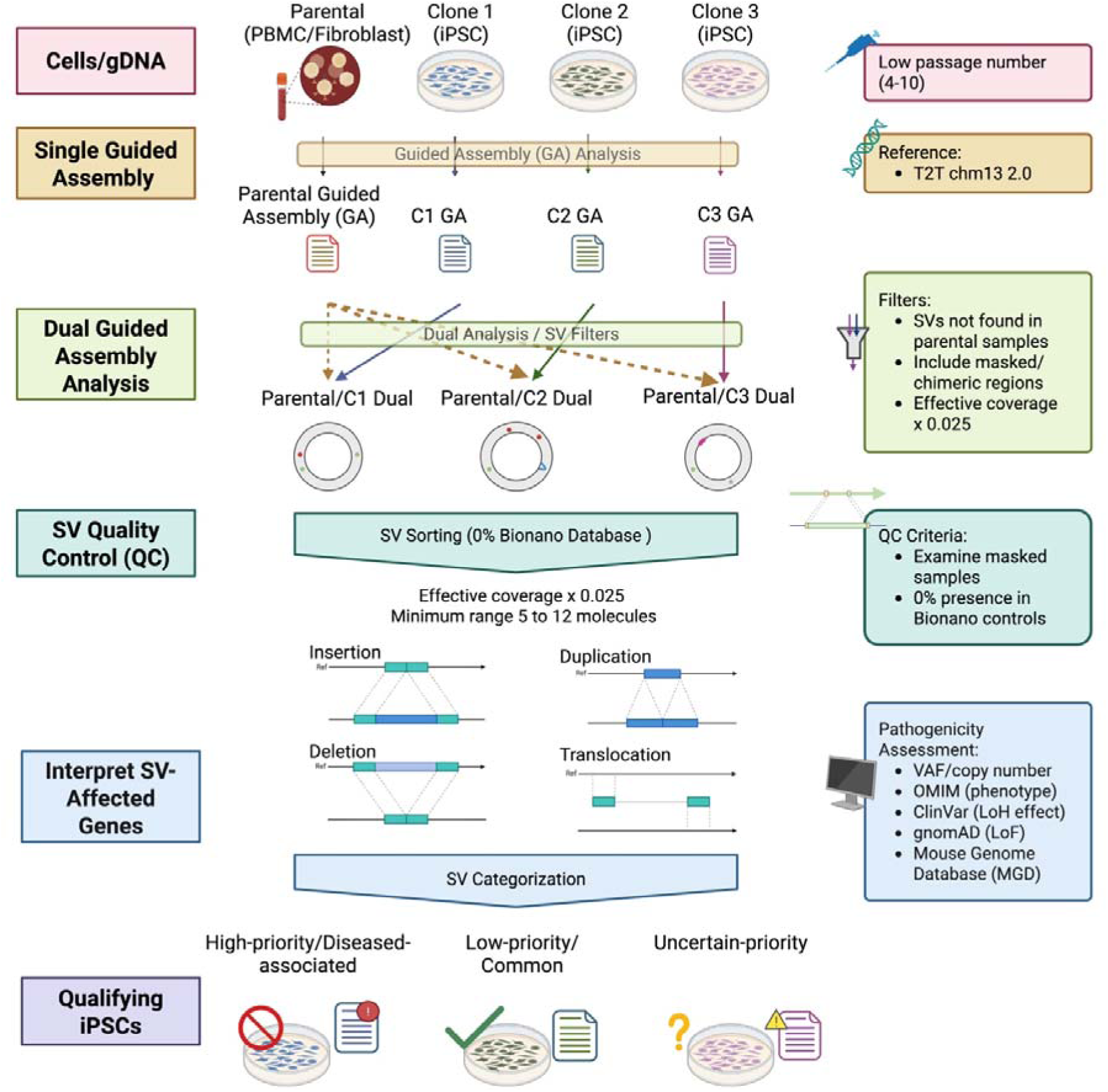
Overview of Analysis Workflow for OGM. After extraction and processing, sample-read molecules were assembled into Guided Analysis SV data, compared between iPSC clones and parental lines (Dual Analysis), then filtered by QC metrics and analyzed for SV categorization.

OGM quality metrics were similar between PBMC-iPSCs and FBC-iPSCs, and all samples satisfied the per-sample QC criteria outlined in **Table S2,** supporting comparability of downstream SV analyses across groups. We applied a QC-focused filtering strategy to prioritize high-confidence, clone-specific events. SVs were retained only if they were not supported above the study-defined threshold in the parental sample, masked/chimeric regions were reviewed as needed, and a sample-specific minimum molecule support threshold was set using effective coverage × 0.025 to capture low-frequency events while limiting background noise, which yielded typically ∼5–12 molecules per variant (**Table S3**). We further prioritized variants that were absent in the Bionano control database [22].

Finally, we classified SVs by type (insertion, deletion, duplication, translocation) and evaluated their potential biological or disease relevance by annotating SV-overlapping genes and summarizing evidence from external resources, including OMIM, ClinVar, gnomAD, constraint metrics, and mouse phenotype databases. The resulting annotations were used to qualify iPSC lines for downstream applications, flagging clones with potentially deleterious SVs and monitoring variants of uncertain priority.

### FBC-iPSCs carry more SVs per clone and present a distinct SV-type profile

Across 39 PBMC-iPSC clones, we identified 69 clone-specific SVs. The SV spectrum was dominated by deletions and insertions: 49% deletions (34/69), 41% insertions (28/69), 7% duplications (5/69), and 3% translocations (2/69) (**Figure 3A, Table S4**). The number and type of SVs varied across donors and sister clones within each set. Overall, PBMC-iPSC clones showed a lower per-clone SV burden (mean 1.86 SVs/clone, median 1, IQR 0–3). Overall, thirteen of 39 clones (33%) showed no detectable SVs (**Table S6**). PBMC-iPSCs were screened for T cell receptor (TCR) chains, and only one SV involving TCRβ (7q35) was identified, indicating that TCR rearrangements do not significantly impact overall SV counts [23].

**Figure 3:**
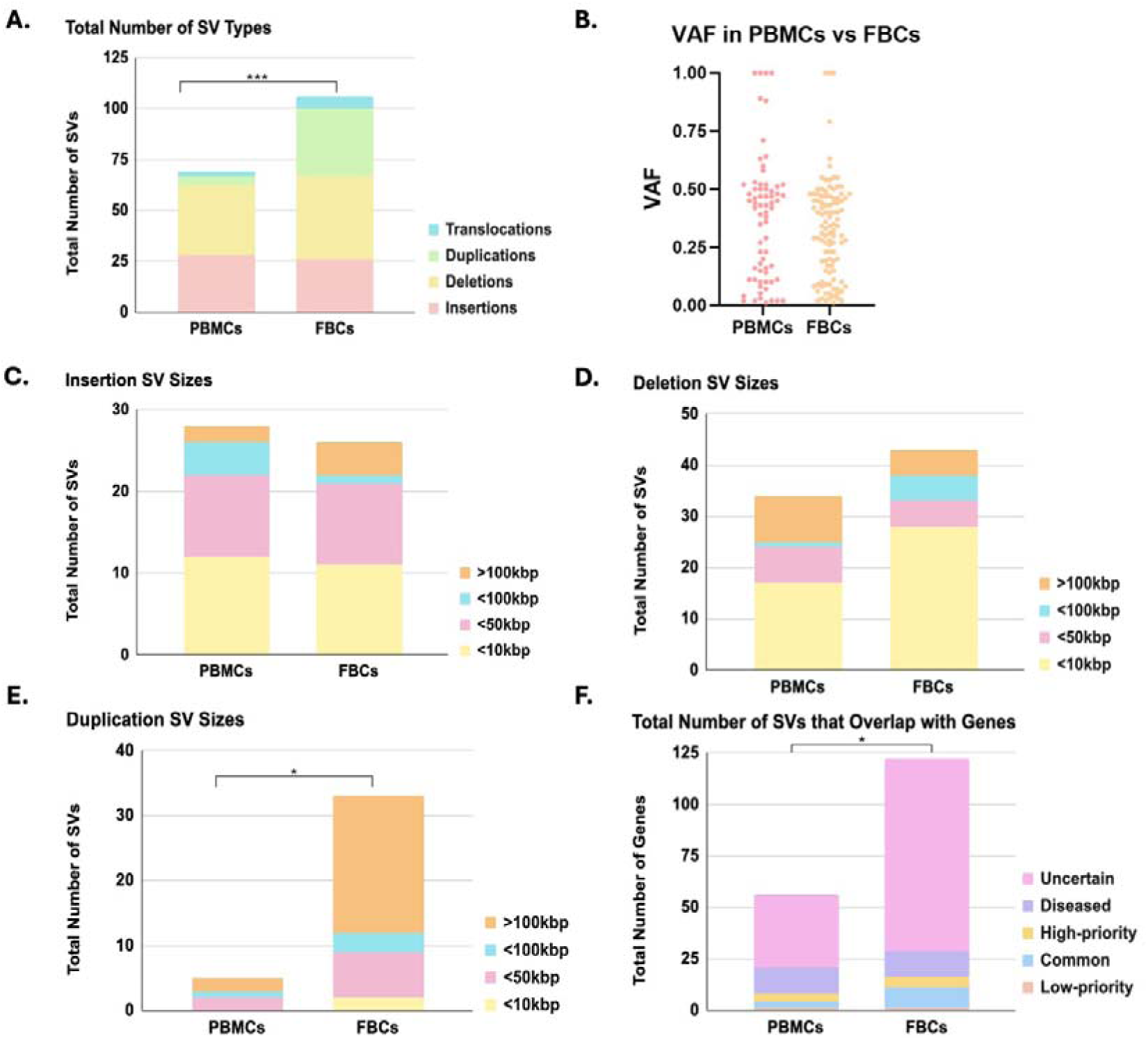
Structural Variant Characteristics and Gene Overlap. A. SV counts differed significantly between groups (2x4 chi-square, ***p = 0.0010). PBMC-iPSCs: 69 SVs (mean 1.86, median 1, IQR 0-3); FBC-iPSCs: 106 SVs (3.12, median 3, IQR1-4). B. VAF range: 0.1–1.0 (median: PBMC-iPSC = 0.42; FBC-iPSC = 0.34). FBC-iPSCs: more low-frequency variants (<0.2; n = 34, median 0.08) than PBMC-iPSCs (n = 24, median 0.10; Mann–Whitney p = 0.862). VAF was mostly heterozygous (PBMC median 0.42 [IQR 0.11–0.51], n = 69; FBC median 0.34 [IQR 0.16–0.47], n = 106; p = 0.22). C. PBMC-iPSCs: 28 insertions across 4 size bins (12, 10, 4, and 2 SVs); FBC-iPSCs: 26 insertions (11, 10, 1, and 4 SVs). Median insertion sizes were similar (PBMC: 11.8 kbp [IQR 4.5–48.0]; FBC: 12.0 kbp [IQR 8.1–48.9]; p = 0.65). D. PBMC-iPSCs: 34 deletions across 4 size bins (17, 7, 1, and 9 SVs); FBC-iPSCs: 41 deletions (27, 4, 5, and 5 SVs). Median deletion sizes differed (PBMC: 14.5 kbp [IQR 3.1–141]; FBC: 5.6 kbp [IQR 3.2–89.4]; p = 0.50). E. PBMC-iPSCs: 5 duplications across 4 size bins (0, 2, 1, and 2 SVs); FBC-iPSCs: 33 duplications (2, 7, 3, and 21 SVs) and larger ≥100kbp (PBMC: median 53.2 kbp [IQR 38.1 kbp–1.06 Mbp], n = 5; FBC: median 203.7 kbp [IQR 66.4–709.8 kbp], n = 33, 2x4 chi-square *p = 0.033; Cramér’s V = 0.462). F. SVs overlapping disease-associated genes were higher in FBC-iPSCs than PBMC-iPSCs (3.53 vs 1.44 genes/clone; IRR = 2.46, 95% CI 1.79–3.37; p = 2.7×10^-8^). PBMC-iPSCs: more high-priority overlap (30.4% vs 15%; OR = 2.47, 95% CI 1.16–5.27; Fisher’s *p = 0.0195). FBC-iPSCs: more uncertain-priority overlap (76.6% vs 62.5%; p = 0.052). Low/common-priority overlap did not differ (p = 1.0). *p < 0.05, ***p < 0.001; n.s.(p > 0.05).

Across 34 FBC-iPSC clones, we identified 106 clone-specific SVs (**Table S5**). Compared with PBMC-iPSCs, the SV spectrum shifted toward deletions and duplications: 39% deletions (41/106), 31% duplications (33/106), 24% insertions (26/106), and 6% translocations (6/106) (**Figure 3A**). As with PBMC-iPSCs, SV burden varied across donors and sister clones. Overall, FBC-iPSC clones showed a higher per-clone SV burden (mean 3.12 SVs/clone, median 3, IQR 1–4). Only five of 34 clones (15%) had no detectable SVs (**Table S7**).

When the total SV burden per clone was compared between groups, FBC-iPSCs showed more SVs. This difference was significant by Mann–Whitney U (p = 0.008), consistent with skewed, over-dispersed count data. SV-type composition also differed when SVs were pooled across clones (2×4 chi-square p ≈ 0.0010), driven by the higher number of duplications in FBC-iPSCs. Finally, PBMC-iPSCs were more likely to be SV-free (33% vs 15%), although this difference did not reach significance by Fisher’s exact test (p = 0.101).

Because breakpoint localization is less reliable in acrocentric p arms and pericentromeric heterochromatic regions, SVs in these regions were excluded from downstream analyses (**Table S8**).

### VAF distributions in PBMC-iPSCs and FBC-iPSCs were centered near the heterozygous range

VAF was compared between PBMC-iPSCs and FBC-iPSCs (**Figure 3B**), with values ranging from 0.08 to 1.0. The median VAF was 0.42 in PBMC-iPSCs and 0.34 in FBC-iPSCs. FBC-iPSCs showed a greater number of low-frequency variants (≤ 0.2; n = 34, median 0.08) compared with PBMC-iPSCs (n = 24, median 0.1), although this difference was not statistically significant (Mann–Whitney p = 0.862). High-frequency variants (≥ 0.75) were observed in both groups but occurred less often than low-frequency events (PBMC n = 6; FBC n = 4). VAF distributions in PBMC-iPSCs (median 0.42 [IQR 0.13–0.51], n = 69) and FBC-iPSCs (median 0.34 [IQR 0.15–0.46], n = 106) were centered near the heterozygous range but includes a substantial subset of low-frequency events. Although FBC-iPSCs showed a higher number of low-frequency variants, the overall difference between groups was not statistically significant (Mann–Whitney p = 0.22).

### Insertions and deletions were of similar size in PBMC- and FBC-iPSCs, but FBC-iPSCs were enriched for larger duplications

OGM can detect SVs ranging from approximately 2 kbp to several Mbp. In PBMC-iPSCs, we found SV sizes from 2,036 bp to 1,735,545 bp, with a median size of 12,027 bp and an average size of 109,032 bp. In FBC-iPSCs, SV sizes ranged from 2,045 bp to 4,905,268 bp, with a median size of 20,535 bp and an average size of 579,760 bp.

We binned SV sizes as <10 kbp, 10–<50 kbp, 50–<100 kbp, and >100 kbp, excluding translocations (size reported as unknown/–1). When SV sizes were compared within each SV class (insertion, deletion, duplication, excluding translocations), PBMC-iPSCs and FBC-iPSCs showed broadly similar size distributions for insertions and deletions, with most events in the low-kbp range. Insertions had nearly identical medians (PBMC: 11.8 kbp [IQR 4.5–48.0], n = 28; FBC: 12.0 kbp [IQR 8.1–48.9], n = 26; Mann–Whitney p = 0.65) (**Figure 3C**). Deletions showed different medians (PBMC: 14.5 kbp [IQR 3.1–141], n = 34; FBC: 5.6 kbp [IQR 3.5–29], n = 41; p = 0.50) (**Figure 3D),** though PBMC included a wider upper tail due to a 1.74 Mb event. Duplications were far more frequent in FBC-iPSCs and tended to be larger (PBMC: median 53.2 kbp [IQR 38.1 kbp–1.06 Mbp], n = 5; FBC: median 203.7 kbp [IQR 66.4–709.8 kbp], n = 33, 2×4 chi-square p = 0.033; Cramér’s V = 0.462) (**Figure 3E)** but the formal comparison was not significant (p = 0.86) and is underpowered for PBMC because there were very few PBMC duplications.

### FBC-iPSCs showed more SV-overlapping protein-coding genes overall, but most gene overlaps remained of uncertain priority

We identified SVs overlapping protein-coding genes in both PBMC-iPSCs and FBC-iPSCs and annotated these genes for prior disease association, neurological relevance, and clone-selection priority. As these iPSC clones were generated for modeling of neurological diseases, we further evaluated whether the affected genes had reported associations with neurodevelopmental, neurological, or neurodegenerative disorders. A total of 56 protein-coding genes were identified in PBMC-iPSCs, with 2% (1/56) classified as low-priority gene overlap, 5% (3/56) as common gene overlap, 7% (4/56) as high-priority disease gene overlap, 23% (13/56) as disease-associated gene overlap, and 63% (35/56) of uncertain priority (**Figure 3F**). Among the 17 high-priority disease and disease-associated gene overlaps, nine were linked to neurodevelopmental disorders, and three were associated with neurodegenerative diseases (**Table S9**).

In FBC-iPSCs, SVs overlapped 120 protein-coding genes, with 1% (1/120) classified as low-priority gene overlap, 8% (10/120) as common gene overlap, 4% (5/120) as high-priority disease gene overlap, 11% (13/120) as disease-associated gene overlap, and 76% (92/120) of uncertain priority (**Figure 3F**). In FBC-iPSCs, of the 13 disease-associated gene overlaps and five high-priority associated gene overlaps, ten were linked to neurodevelopmental disorders, one to a neurodegenerative disease, and the remaining seven were not associated with neurological conditions (**Table S10**). When comparing PBMC-iPSCs and FBC-iPSCs, a greater number of protein-coding genes were identified in FBC-iPSCs (p = 0.0269). In comparison, PBMC-iPSCs were associated with a higher number of genes linked to neurodegenerative diseases compared to FBC-iPSCs.

Normalizing by the number of clones, FBC-iPSCs showed a ∼2.46× higher rate of affected genes per clone (3.53 vs 1.44 genes/clone; IRR = 2.46, 95% CI 1.79–3.37; p = 2.7×10⁻^8^, Poisson rate comparison). In PBMC-iPSCs, 17/56 genes (30.4%) fell in the high-priority/disease-associated categories versus 18/120 (15%) in FBC-iPSCs (OR = 2.47, 95% CI 1.16–5.27; Fisher’s p = 0.0195), indicating a higher higher proportion of high-priority or disease-associated gene overlaps among PBMC-iPSC gene hits despite fewer total genes affected. FBC-iPSCs also showed a larger share of variants of uncertain priority (76.6% vs 62.5%; Fisher’s p = 0.052), while low-priority /common gene overlap proportions did not differ (Fisher’s p = 1.0).

### A small fraction of SVs overlapped recurrent hPSC culture-associated chromosomal regions

SV analysis identified insertions, deletions, and duplications that mapped to chromosomal regions repeatedly reported as cell culture-associated “hotspots” in human pluripotent stem cells (hPSCs), including intervals on chromosomes 1, 12, 17, 18, and 20[24][25]When we mapped SVs onto recurrent hPSC culture–associated regions reported by Baker et al. 2016[24] (e.g., 20q11.21, 12p11–pter, 1q gains, and recurrent 17q/18q events), overall, only a small fraction of SVs overlapped these regions. In PBMC-iPSCs, four hotspot-region SVs were identified, two at 12p11.21 and two at 20q11.21 out of 69 total SVs (**Table 1**). FBC-iPSCs exhibited seven SVs, one at 1q25.1, two within 1q31.1-1q36.13, two within 12p11.1–p11.21, one at 17q25.3, and one at 18q23 out of 106 total SVs (**Table 2**)[24]. The chromosome location 1q25.1 overlaps with the fragile site mentioned in **Table 4**.

**Table 1:** Recurrent hPSC culture-associated chromosomal regions overlapped by four PBMC-iPSC SVs.

| SUSL Line | Chromosome | Size | SV Type | SV-overlapping Genes | Description |
| --- | --- | --- | --- | --- | --- |
| SUSL 63 C6 | 12p11.21 | 43,453 | Insertion | DENND5B | DENND5B is associated with neurodevelopmental phenotypes featuring cognitive impairment, dysmorphism, abnormal behavior, variable epilepsy, white matter abnormalities, and cortical gyration defects [26]. |
| SUSL 63 C6 | 12p11.21 | 457,364 | Deletion | BICD1, EFGD4 | BICD1 is associated with hearing loss and peripheral neuropathy [27]. FGD4 is associated with Charcot-Marie-Tooth disease, type 4H[28]. |
| SUSL 88 C7 | 20q11.21 | 130,523 | Deletion | - | Chromosome 20q11.21 associated with feeding problems, retinal dysplasia, and skeletal abnormalities [29]. |
| SUSL 63 C6 | 20q11.21 | 4,488 | Insertion | - | Chromosome 20q11.21 associated with developmental delays and cancer generation [30] |

**Table 2:** Recurrent hPSC culture-associated chromosomal regions overlapped by seven FBC-iPSC SVs.

| SUSL Line | Chromosome | Size | SV Type | SV-overlapping Genes | Description |
| --- | --- | --- | --- | --- | --- |
| SUSL 39 C4 | 1q25.1 | 203,673 | Duplication | RC3H1, RABGAP1L | RC3H1 is associated with immune dysregulation and systemic hyperinflammation syndrome [31]. |
| SUSL 07 C5 | 1q31.1 | 10,557 | Deletion | - | Chromosome 1q31.1 is associated with congenital heart disease [32]. |
| SUSL 45 C3 | 1q36.13 | 228,825 | Deletion | - | Chromosome 1q36.13 is associated with learning disabilities, behavioral abnormalities, and ptosis [33]. |
| SUSL 07 C9 | 12p11.1 | 19,371 | Insertion | - | Chromosome 12p11.1 is associated with Noonan syndrome [34]. |
| SUSL 73 C10 | 12p11.21 | 3,434 | Insertion | - | Chromosome 12p11.21 is associated with Noonan syndrome [34]. |
| SUSL 66 C4 | 17q25.3 | 3,703 | Deletion | METRNL | METRNL is associated with type 2 diabetes, obesity, coronary heart disease and atherosclerosis, colitis, psoriasis and arthritis [35]. |
| SUSL 39 C4 | 18q23 | 5,158 | Deletion | NFATC1 | Chromosome 18q23 and NFATC1 associated with head and neck squamous carcinoma [36]. |

Overall, these findings indicate that while a subset of iPSC clones carried SVs within recurrent hPSC culture-associated chromosomal regions, most clone-specific SVs occurred outside these well-described recurrent loci[37].

### Fragile-site SVs were rare overall in PBMC-iPSCs and FBC-iPSCs

PBMC-iPSCs contained three SVs overlapping annotated fragile sites, consisting of one insertion and two deletions on chromosomes 2, 3, and 9 (**Table 3).**

**Table 3:**
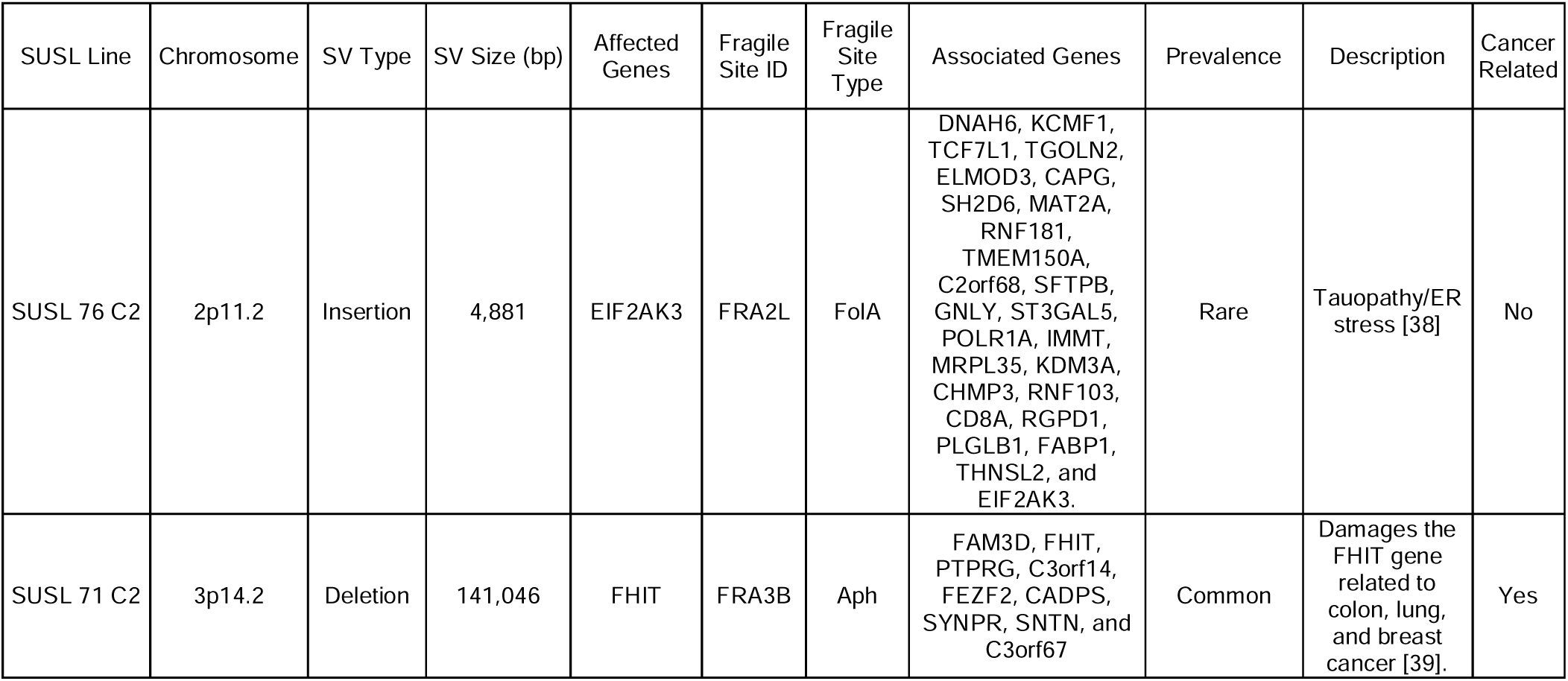
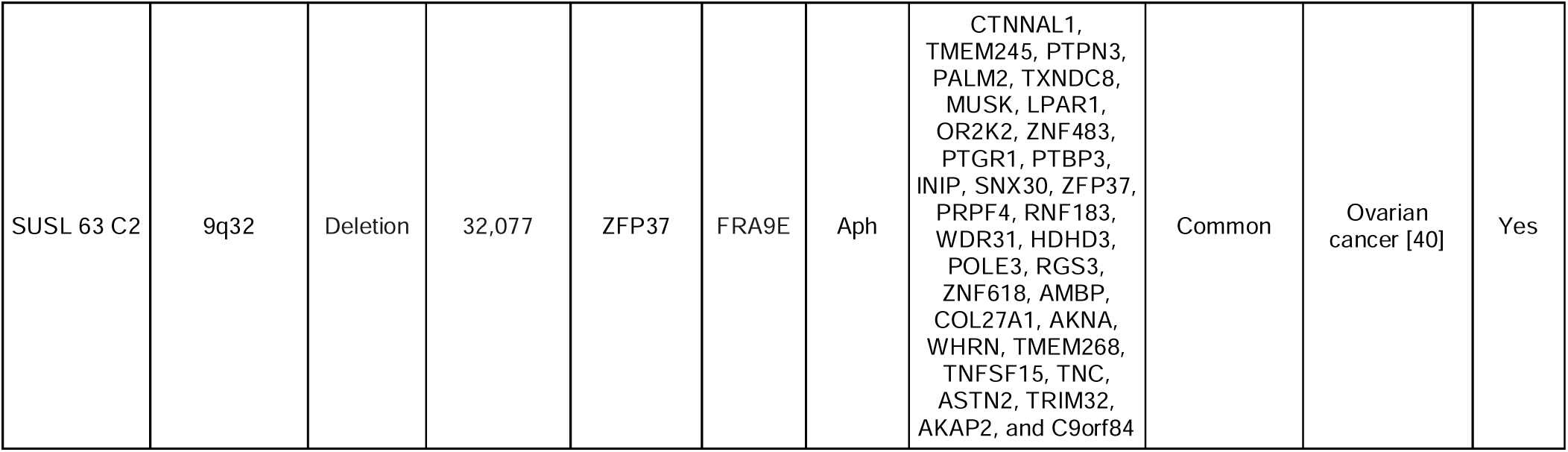
3 fragile sites (FRA2L, FRA3B, & FRA9E) identified in 3 PBMC-iPSCs.

The two deletions overlapped the common fragile sites FRA3B (3p14.2) and FRA9E (9q32), intersecting the *FHIT* and *ZFP37* genes, and these aphidicolin-sensitive regions have been associated with cancer [39][40]. The single insertion overlapped the fragile site FRA2L (2p11.2), which is considered rare, and occurs within the *EIF2AK3* gene.

In comparison, FBC iPSCs showed five SVs overlapping annotated fragile sites, two duplications, and three deletions across chromosomes 1, 2, and 14 (**Table 4**). One duplication overlapped the common aphidicolin-sensitive fragile site FRA1G (1q25.1). The second duplication overlapped the rare folate-sensitive fragile site FRA2K (2q22.3), intersecting the *ACVR2A* gene.

**Table 4:**
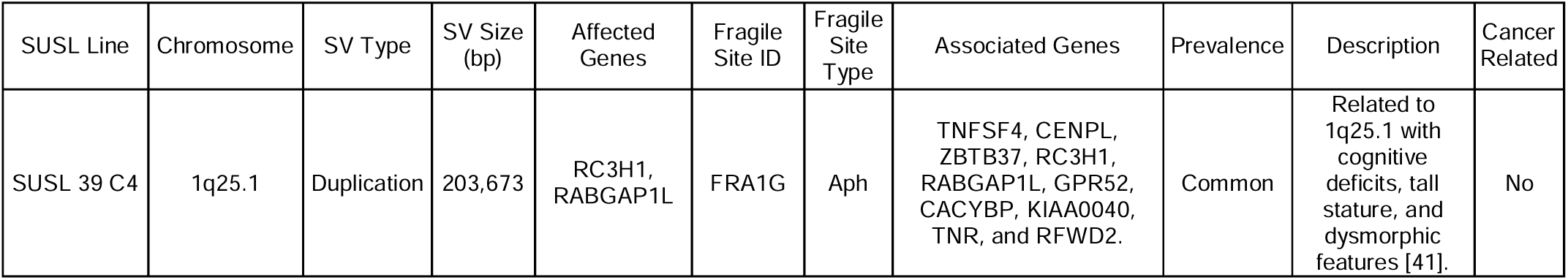
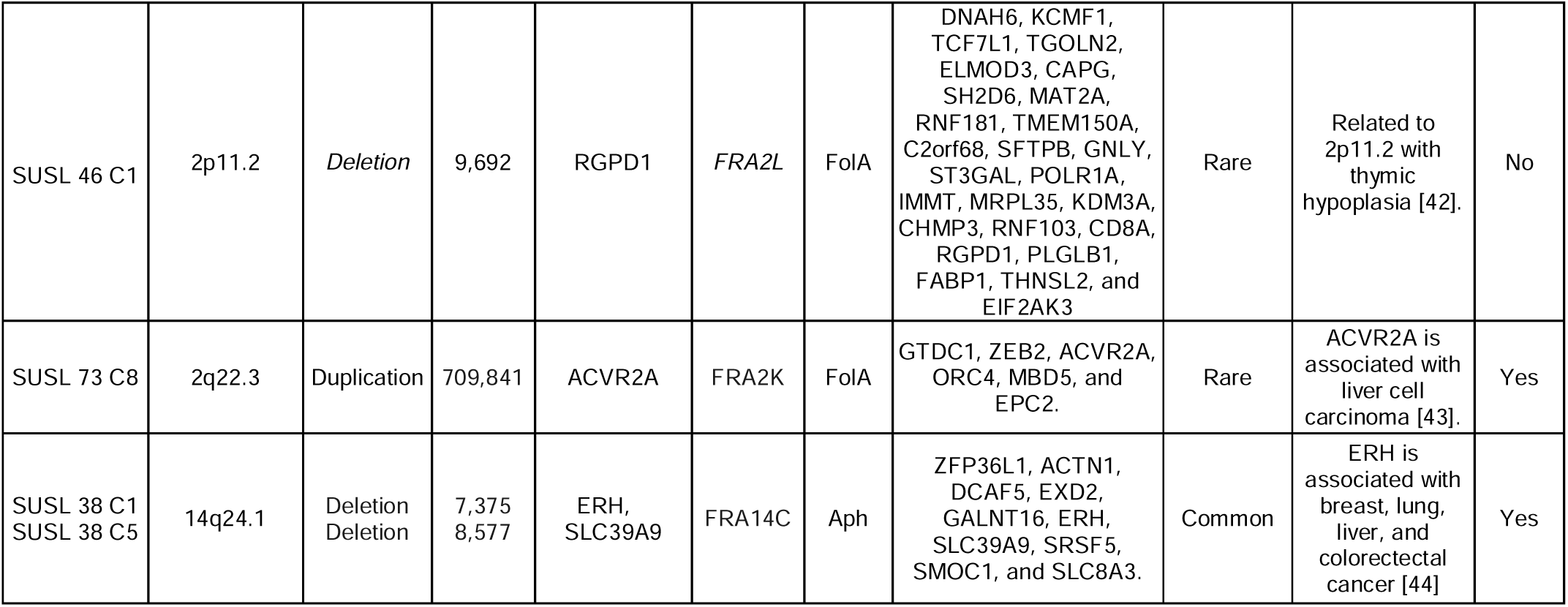
4 fragile sites (FRA1G, FRA2L, FRA2K, & FRA14C) identified in 5 FBC-iPSCs.

One deletion overlapped the rare fragile site FRA2L and intersected *RGPD1*, rather than *EIF2AK3*, as observed in PBMCs. Two additional deletions overlapped the common fragile site FRA14C and intersected ERH and SLC39A9.

Overall, fragile-site overlap accounted for only a small subset of all SVs in both groups. PBMC-iPSCs showed fewer fragile-site SVs overall, including events at two well-characterized cancer-associated common fragile sites, whereas FBC-iPSCs showed a slightly higher number of fragile-site SVs distributed across both common and rare fragile-site classes.

### Acrocentric p-arm SV calls were common but hard to interpret for iPSC QC

We also detected SV calls on acrocentric short arms, including 13p, 14p, 15p, 21p, and 22p, in both PBMC- and FBC-derived iPSC clones. In total, 38 acrocentric p-arm SV calls were identified, including 25 in PBMC-iPSCs and 13 in FBC-iPSCs (**Table S8**). Because acrocentric p-arms and pericentromeric regions are highly repetitive (e.g., rDNA and satellite arrays), these calls are difficult to interpret by OGM and may reflect repeat-length variation, reduced alignment specificity, or reference-mapping uncertainty rather than culture-acquired, gene-disrupting SVs. For this reason, acrocentric p-arm SV calls were reported separately and excluded from downstream burden, gene-overlap, hotspot, and fragile-site analyses.

## DISCUSSION

### Genomic integrity remains a central constraint for disease-model iPSCs

iPSCs enable scalable, patient-relevant models of human disease, but reprogramming and culture can introduce or select genomic changes that may confound downstream phenotypes[45][46][47][48][49][50][5]. Large-scale genomic studies of human iPSCs have shown that human iPSCs frequently acquire de novo copy number and structural variants during reprogramming and early culture, resulting in strong clonal selection shaping the genomic landscape. Previous studies suggest that SVs arise directly as a consequence of the reprogramming process or from subclonal mutations in the parental cell population that become enriched during reprogramming and expansion. Both mechanisms may contribute, with some SVs emerging in response to the cellular stress associated with reprogramming, while others may reflect pre-existing mosaic variants carried over from the starting cells. However, evidence suggests that reprogramming and subsequent culture can act as key drivers of genomic instability in iPSCs[51][52][53][54]. SVs are particularly important because they can disrupt genes and regulatory landscapes and because low-frequency or balanced events can evade routine screening[55][30][56][57] Many SVs, particularly mosaic or balanced events, are difficult to detect using standard quality-control methods such as karyotyping or CNV microarrays. The presence of heterozygous and lower-frequency SVs in our clones is therefore consistent with prior studies showing that reprogramming and early clonal expansion can generate or enrich clone-specific genomic changes[58][59]. Although some mosaic variants may originate from the parental cell population and remain below the detection threshold in bulk parental samples, our paired OGM approach allowed us to prioritize SVs that were not supported above the study-defined threshold in the matched parental line. Specifically, we used sample-specific molecule-support thresholds based on effective coverage, retaining heterozygous or mosaic SV calls when supported by effective coverage × 0.025, which typically corresponded to approximately 5–12 supporting molecules per variant. This filtering strategy enabled the detection of clone-enriched SVs, including lower-frequency events, while reducing background calls.

### OGM strengthened post-reprogramming QC beyond standard cytogenetic screens

Prior work has emphasized that karyotyping and array-based methods reliably detect large, unbalanced changes but often miss smaller and/or balanced SVs and low–allele fraction events that can influence clone selection[55][30][56][57]. In our initial proof-of-concept analysis of the reference iPSC line KOLF2.1J[60], OGM detected 36 SVs below ∼100 kb that were not reported by prior SNP array–based CNV analysis, which captured only five variants ≥100 kb, suggesting that a substantial fraction of iPSC SV burden falls below the effective resolution of common QC assays. Building on that test-bed, the present study scaled OGM to a cohort of clonal iPSC lines derived from PBMCs and FBCs, where it served as a post-reprogramming QC layer to capture insertions, deletions, duplications, and translocations genome-wide, enable clone-to-parent comparisons, and support downstream annotation for gene overlap, fragile sites, and recurrent regions associated with pluripotent cell adaptation[61][5]. Although implementation will depend on local cost, throughput, and analysis infrastructure, OGM is operationally well-suited as a complementary QC approach when high-resolution SV detection is important for iPSC clone selection.[30][57]

### Somatic cell source was associated with SV burden and SV-type composition

A key observation in our dataset is that FBC-iPSCs showed a higher number of SVs compared to PBMC-iPSCs. Beyond total counts, pooled SV-type composition differed strongly, driven mainly by an enrichment of larger ≥100kbp duplications in FBC-iPSCs. PBMC-iPSCs showed a lower number of SVs, and a greater proportion of PBMC-iPSC clones were SV-free. These observations are consistent with previous reports describing differences in cytogenetic stability between PBMCs and FBCs during reprogramming[62][63][64].

### Recurrent hotspots and SVs at fragile sites appeared, but they represented a small fraction of the total SVs

Recurrent hotspot events appeared in both PBMC-iPSCs and FBC-iPSCs, mapping to chromosomal regions repeatedly reported as prone to culture-acquired change in pluripotent stem cells (e.g., loci on chromosomes 1, 12, 17, 18, and 20) and often interpreted as adaptive gains or losses that can provide a growth or survival advantage during expansion[25][65][66][24][67][68].

However, these hotspot-consistent calls represented only a small fraction of the total SV burden; most SVs fell outside canonical recurrent intervals, supporting a model in which diffuse clone-specific SV detection occurs after reprogramming and early expansion, although low-level parental mosaicism and clonal selection cannot be excluded.

Fragile sites provide a complementary genomic context for interpreting SV localization because these regions are prone to replication stress, DNA breakage, and rearrangement. In our dataset, SVs overlapped both common and rare fragile sites, with a slightly higher number of fragile-site overlaps in FBC-iPSCs than in PBMC-iPSCs. This pattern aligns with the idea that somatic starting populations differ in replication history or susceptibility to reprogramming-associated stress, which may contribute to differences in the distribution of clone-specific SVs after reprogramming and early expansion [19][69][63][64].

### Gene-overlapping SVs underscored the need for clone-level interpretation

We observed gene-overlapping SVs in iPSC clones from both cell sources, including variants annotated as high-priority/disease-associated gene overlap, but also many variants of uncertain priority. This pattern matches the broader iPSC literature: structural changes often land in or near genes, but functional relevance frequently requires orthogonal evidence[47][49][69]. Because iPSCs are clonal products of reprogramming and early expansion, clone-to-clone variability can persist even within a donor, reinforcing the need for clone-level genomic QC and thoughtful line selection for disease modeling[51] [70][69][5].

### Technical and experimental limitations

This study has several limitations. First, we could not directly compare both PBMC- and fibroblast-derived iPSC clones generated from the same donor because paired starting materials were not available for all individuals. Therefore, the observed differences between groups should be interpreted as source-associated rather than strictly source-causal, as donor biology, diagnosis, reprogramming method, provider, passage history, and culture conditions may also contribute to the observed SV profiles. Future studies using matched PBMC and fibroblast samples from the same donors will be useful to further separate the contribution of somatic cell source from donor-specific and technical factors.

Second, this study was designed as an OGM-based clone-parent comparison rather than a direct benchmarking study against conventional cytogenetic assays. Same-sample karyotyping, FISH, or array-based analyses were therefore not performed for all clones. However, the paired OGM design enabled high-resolution detection of clone-specific SVs and provided a practical framework for post-reprogramming genomic QC.

Third, SV detection in repetitive or structurally complex regions remains challenging; we recorded but excluded acrocentric p-arm calls from downstream analysis due to alignment constraints. In addition, low-level mosaic variants inparental somatic populations may fall below the detection threshold and could be enriched during clone selection, complicating the interpretation of variants that appear clone-specific [5][64].

Finally, clone selection during iPSC derivation may generate sister clones with shared early events, and the number of clones per donor limits donor-stratified inference. Larger multi-site studies with harmonized reprogramming methods, matched somatic sources, and standardized passage histories will help clarify how starting material, donor background, and culture conditions shape SV burden in iPSC clones [63] [64][71].

## Conclusions

This study demonstrates the utility of OGM as a high-resolution, clone-level genomic QC approach for early-passage iPSC lines. By comparing each iPSC clone with its matched parental somatic line, OGM enabled detection and annotation of clone-specific SVs at approximately ≥2 kbp, including events overlapping protein-coding genes, fragile sites, and recurrent hPSC culture-associated chromosomal regions. In this cohort, fibroblast-derived iPSC clones showed a higher SV burden, including more frequent and larger duplications, whereas PBMC-derived iPSC clones were more often free of detectable clone-specific SVs. These findings support the use of OGM as a complementary post-reprogramming QC strategy and suggest that somatic cell source may be associated with distinct SV profiles in disease-model iPSC clones.

## Abbreviations

ADRC: Alzheimer’s Disease Research Center
AD: Alzheimer’s disease
CNV: Copy number variation
FBC: Fibroblast cells
hPSCs: Human pluripotent stem cells
iPSCs: Induced pluripotent stem cells
LRS: Long-read sequencing
NGS: Next-generation sequencing
OGM: Optical genome mapping
PBMC: Peripheral blood mononuclear cells
PD: Parkinson’s disease
QC: Quality control
SV: Structural variant
TCR: T cell receptor
UHMW: Ultra-high molecular weight
VAF: Variant allele frequency
WGS: Whole-genome sequencing

## Declarations

### Ethics approval and consent to participate

Cell lines were derived from donors who gave written consent under Stanford University IRB 33727 and IRB 48895. The iPSC lines are covered under Stanford University Stem Cell Research Oversight (SCRO) protocol 567 and 754.

### Consent for publication

Not applicable

### Availability of data and materials

The data have been deposited in the European Nucleotide Archive (ENA) under project accession number PRJEB91134 and (Secondary accession number: ERP174125) under the file types BNX, CMAP, SMAP, and XMAP.

### Authors’ contributions

K.S. and F.Z. cultured the reprogrammed cells and collected cell pellets for Bionano analysis. M.J.Y. designed the OGM pipeline and workflow. M.J.Y. and K.S. extracted and labeled gDNA and loaded the samples onto the Saphyr flow cell chip. M.J.Y. initiated the data analysis, which was completed by L.N. L.N. wrote the first draft of the manuscript. R.H. uploaded the raw data files to ENA and cross-checked the ENA database. B.S. conceived the study, provided oversight, reviewed the data analysis, and obtained funding for the study. All authors reviewed and approved the final draft.

## Acknowledgements

Bu Wei He supported the ENA upload of raw data and curated disease-associated gene annotations. Dr. Vittorio Sebastiano generated several iPSC clones.

---

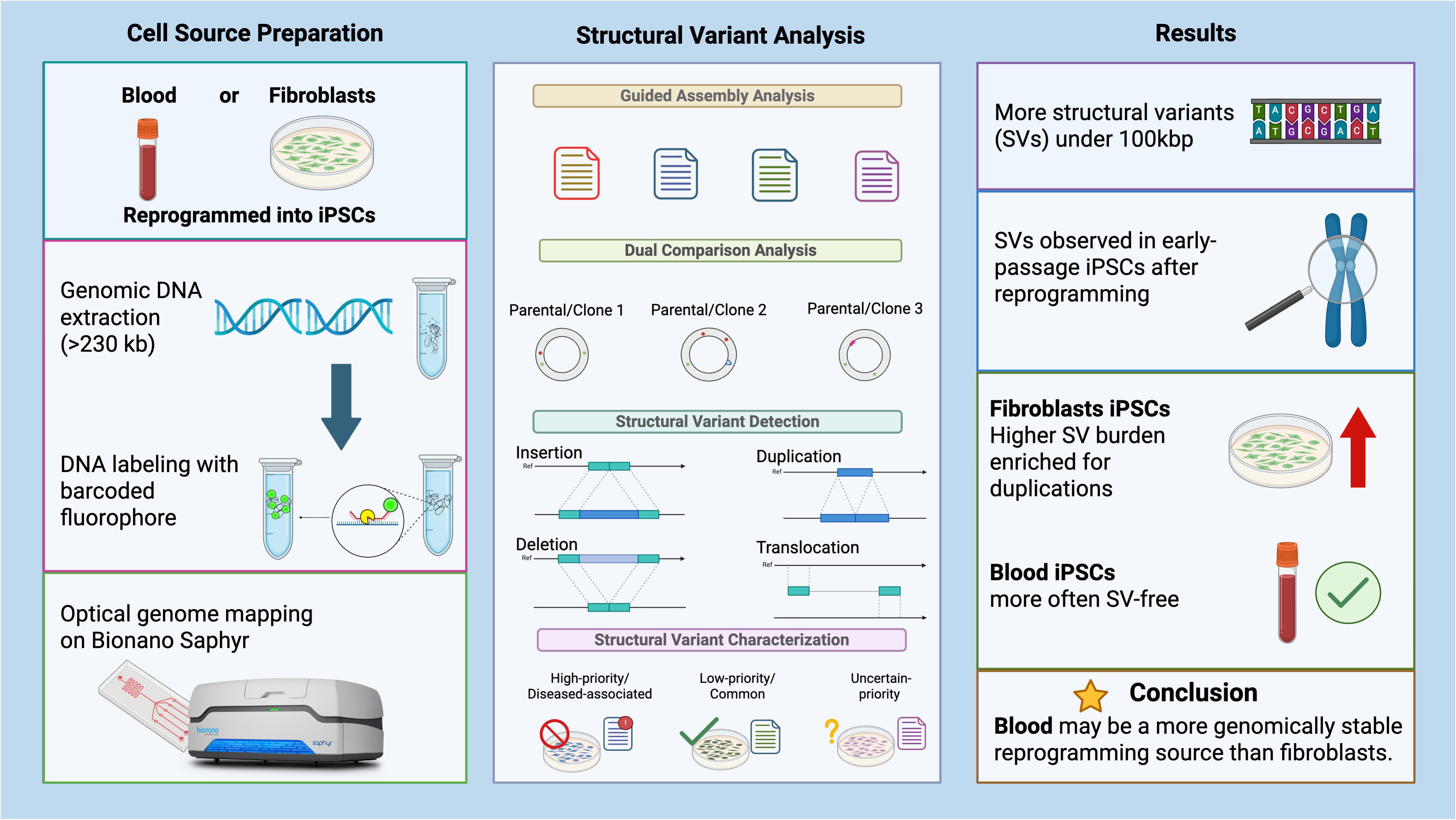

