## Supplemental Figures/Tables for "Optical genome mapping identifies source-associated structural variant differences across early-passage human iPSCs"

**Supplemental Figures and Tables**

**Tables S4, S5, S9, and S10 are in an Excel spreadsheet**

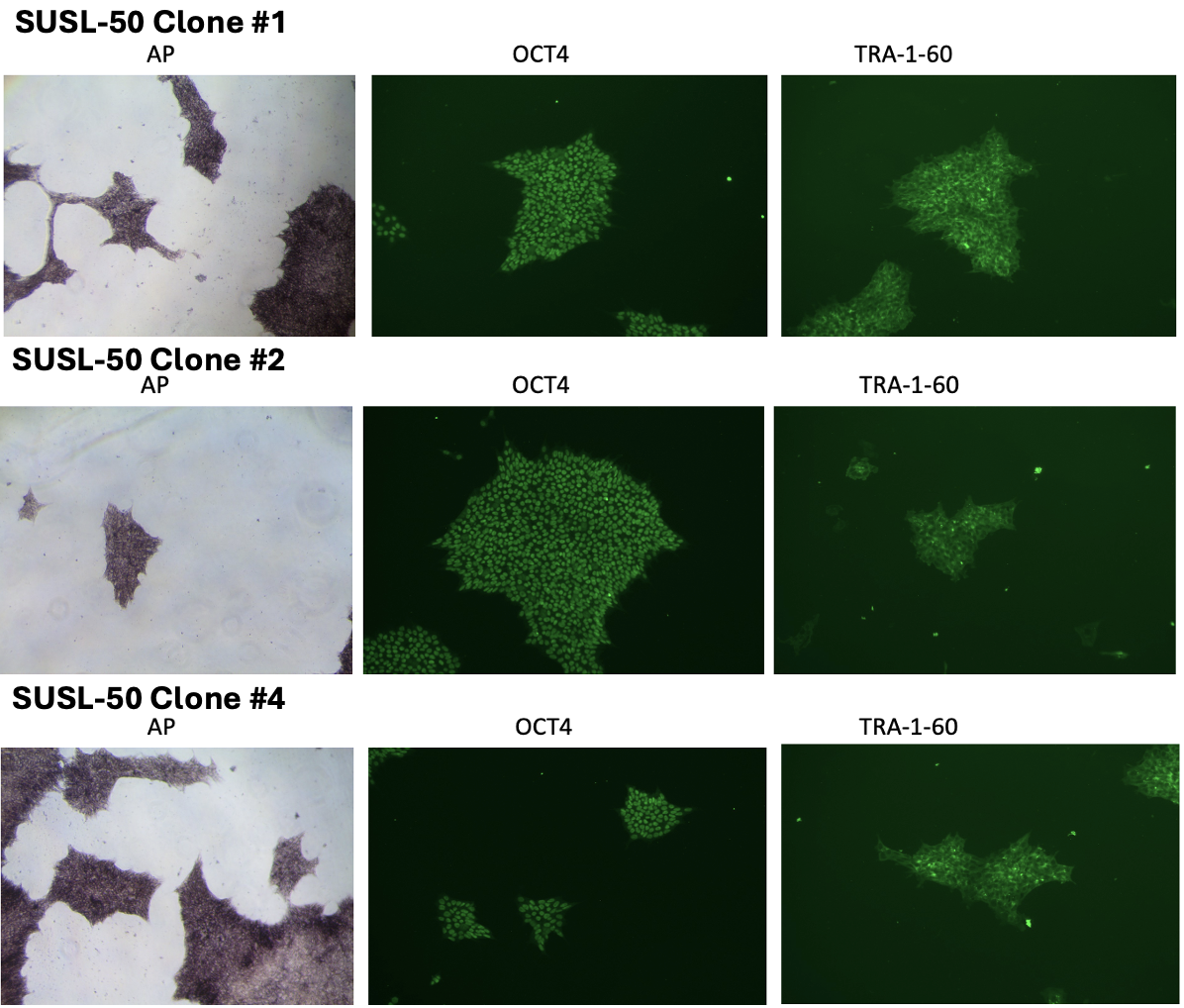

**Figure S1: Pluripotency Characterization of PBMC-derived iPSC Clones.** As an example, SUSL-50 (PBMC donor) was reprogrammed into iPSCs using non-integrating episomal vectors, resulting in transgene- and virus-free clones. Pluripotency was confirmed across all clones using standard characterization assays, including immunostaining for TRA-1-60 and OCT4, as well as assessment of alkaline phosphatase (AP) activity. Pluripotency evaluation was performed for all PBMC-derived iPSC clones.

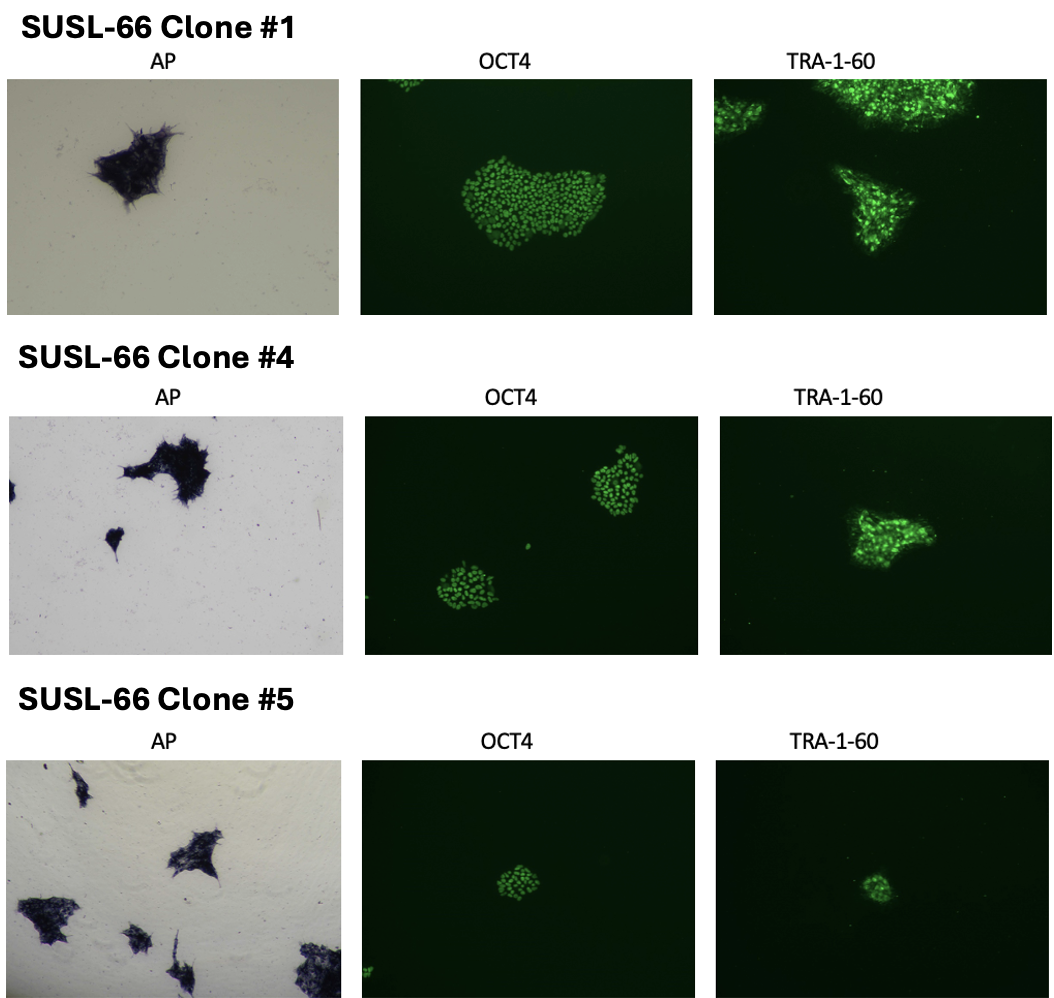

**Figure S2: Pluripotency Characterization of FBC-derived iPSC Clones.** As an example, SUSL- 66 (FBC donor) was reprogrammed into iPSCs using non-integrating, episomal vectors, resulting in transgene- and virus-free clones. Pluripotency was confirmed across all clones using standard characterization assays, including immunostaining for TRA-1-60 and OCT4, as well as assessment of alkaline phosphatase (AP) activity. Pluripotency evaluation was performed for all FBC-derived iPSC clones.

**Table S1: Demographics and Cell Reprogramming Methodology for 25 Donors**

| **Cell Type** | **SUSL Line** | **Sex** | **Age** | **Disease** | **Genotype** | **Reprogramming Method** | **Reprogramming Service** | **Clones** | **Passage Number** |
| --- | --- | --- | --- | --- | --- | --- | --- | --- | --- |
| PBMC | SUSL 50 | M | 70 | Cognitively Unimpaired | E4/E4 | Episomal | ALSTEM  Cell Advancements | SUSL 50 C1  SUSL 50 C7  SUSL 50 C8 | P8  P9  P8 |
| PBMC | SUSL 51 | M | 74 | Cognitively Unimpaired | E4/E4 | Episomal | ALSTEM  Cell Advancements | SUSL 51 C1  SUSL 51 C2  SUSL 51 C4 | P13  P13  P13 |
| PBMC | SUSL 62 | M | 64 | Cognitively Unimpaired | E4/E4 | Episomal | ALSTEM  Cell Advancements | SUSL 62 C1  SUSL 62 C2  SUSL 62 C3 | P8  P7  P7 |
| PBMC | SUSL 63 | F | 74 | Healthy Control | E3/E3 | Episomal | ALSTEM  Cell Advancements | SUSL 63 C1  SUSL 63 C2  SUSL 63 C6 | P8  P10  P8 |
| PBMC | SUSL 64 | M | 72 | Healthy Control | E3/E3 | Episomal | ALSTEM  Cell Advancements | SUSL 64 C1  SUSL 64 C2  SUSL 64 C4 | P7  P6  P7 |
| PBMC | SUSL 70 | F | 60 | Alzheimer's | E4/E4 | Episomal | ALSTEM  Cell Advancements | SUSL 70 C1  SUSL 70 C2  SUSL 70 C3 | P5  P5  P5 |
| PBMC | SUSL 71 | M | 55 | Alzheimer's | E4/E4 | Episomal | ALSTEM  Cell Advancements | SUSL 71 C1  SUSL 71 C2  SUSL 71 C3 | P5  P5  P4 |
| PBMC | SUSL 76 | F | 61 | Alzheimer's | E4/E4 | Episomal | ALSTEM  Cell Advancements | SUSL 76 C1  SUSL 76 C2  SUSL 76 C3 | P5  P5  P5 |
| PBMC | SUSL 77 | M | 75 | Alzheimer's | E4/E4 | Episomal | ALSTEM  Cell Advancements | SUSL 77 C2  SUSL 77 C3  SUSL 77 C4 | P4  P4  P4 |
| PBMC | SUSL 78 | F | 68 | Healthy Control | E3/E3 | Episomal | ALSTEM  Cell Advancements | SUSL 78 C6  SUSL 78 C10  SUSL 78 C11 | P4  P4  P4 |
| PBMC | SUSL 79 | M | 77 | Healthy Control | E3/E3 | Episomal | ALSTEM  Cell Advancements | SUSL 79 C1  SUSL 79 C2  SUSL 79 C4 | P4  P4  P4 |
| PBMC | SUSL 88 | F | 75 | Cognitively Unimpaired | E4/E4 | Episomal | ALSTEM  Cell Advancements | SUSL 88 C2  SUSL 88 C5  SUSL 88 C7 | P8  P8  P8 |
| PBMC | SUSL 90 | F | 76 | Cognitively Unimpaired | E4/E4 | Episomal | ALSTEM  Cell Advancements | SUSL 90 C3  SUSL 90 C6  SUSL 90 C9 | P4  P4  P4 |
| FBC | SUSL 06 | F | 39 | Cognitively Unimpaired | ATXN10 | mRNA | Sebastiano Lab  Stanford | SUSL 06 FBC  SUSL 06 C1  SUSL 06 C2  SUSL 06 C3 | P7  P5  P6  P6 |
| FBC | SUSL 07 | F | 65 | ATXN10 | ATXN10 | mRNA | Sebastiano Lab  Stanford | SUSL 07 FBC  SUSL 07 C3  SUSL 07 C5  SUSL 07 C9 | P7  P4  P4  P4 |
| FBC | SUSL 12 | M | 70 | ATXN10 | ATXN10 | mRNA | Sebastiano Lab  Stanford | SUSL 12 FBC  SUSL 12 A2  SUSL 12 B3  SUSL 12 C1 | P7  P5  P5  P5 |
| FBC | SUSL 13 | F | 38 | ATXN10 | ATXN10 | mRNA | Sebastiano Lab  Stanford | SUSL 13 FBC  SUSL 13 C1  SUSL 13 C2  SUSL 13 C3 | P7  P5  P5  P5 |
| FBC | SUSL 25 | F | 70 | Parkinson’s | G2019S++ | Episomal | Pathways to Stem Cell Science | SUSL 25 FBC  SUSL 25 C2  SUSL 25 C6 | P6  P17  P20 |
| FBC | SUSL 38 | M | 55 | Cognitively Unimpaired | SNCA Duplication | Episomal | ALSTEM  Cell Advancements | SUSL 38 FBC  SUSL 38 C1  SUSL 38 C2  SUSL 38 C5 | P3  P6  P6  P6 |
| FBC | SUSL 39 | F | 54 | Parkinson’s | SCNA Duplication | Episomal | ALSTEM  Cell Advancements | SUSL 39 FBC  SUSL 39 C2  SUSL 39 C4  SUSL 39 C8 | P3  P5  P5  P5 |
| FBC | SUSL 40 | M | 30 | Healthy Control | E3/E3 | Episomal | ALSTEM  Cell Advancements | SUSL 40 FBC  SUSL 40 C1  SUSL 40 C7  SUSL 40 C8 | P4  P4  P4  P4 |
| FBC | SUSL 45 | M | 54 | Cognitively Unimpaired | P62 | Episomal | ALSTEM  Cell Advancements | SUSL 45 FBC  SUSL 45 C3  SUSL 45 C6 | P6  P7  P7 |
| FBC | SUSL 46 | M | 65 | Parkinson’s | G2019S++ | Episomal | ALSTEM  Cell Advancements | SUSL 46 FBC  SUSL 46 C1  SUSL 46 C2  SUSL 46 C9 | P3  P5  P5  P5 |
| FBC | SUSL 66 | M | 74 | Parkinson’s | G2019S-+ | Episomal | ALSTEM  Cell Advancements | SUSL 66 FBC  SUSL 66 C2  SUSL 66 C4  SUSL 66 C5 | P5  P8  P9  P6 |
| FBC | SUSL 73 | F | 55 | Parkinson’s | G2019S-+ PARKIN | Episomal | ALSTEM  Cell Advancements | SUSL 73 FBC  SUSL 73 C3  SUSL 73 C8  SUSL 73 C10 | P6  P5  P5  P5 |

**Table S2: Quality Control gDNA Metrics Thresholds Before and After Labeling**

| **QC Metric** | **Post-Elute gDNA**  **Conc.** | **Post-Label gDNA Conc.** | **CV**  **(σ/ X̅) of Conc.** | **N50> 150kbp** | **N50> 20kbp** | **Coverage** | **Map Rate** | **Label Density** |
| --- | --- | --- | --- | --- | --- | --- | --- | --- |
| **Ideal Range** | 40-150 ng/μL | 4-16 ng/μL | <0.3 | >230kbp | >150kbp | >350X | >80% | 14-17 labels/kbp |
| **Absolute Min.**  **Cutoff** | >40 ng/μL | >3.5 ng/μL | <0.3 | >210kbp | >130kbp | >300X | >70% | >14 labels/kbp |

**Table S3: Bionano SV Filters Used in Workflow**

| **SV Filter Category** | **Filter Name** | **Valid Values** |
| --- | --- | --- |
| **Filter by SV Type** | Insertion Confidence / Minimum Size | Default confidence (0), no minimum |
|  | Deletion Confidence / Minimum Size | Default confidence (0), no minimum |
|  | Inversion Confidence / Minimum Size | Default confidence (0.7), no minimum |
|  | Duplication Confidence / Minimum Size | Default confidence (-1), no minimum |
|  | Intra-Fusion Confidence / Minimum Size | Default confidence (0.02), no minimum |
|  | Inter-Translocation Confidence / Minimum Size | Default confidence (0.02), no minimum |
| **General SV Filters** | Chromosomes To Display | All |
|  | SV Masking Filter | Both “Masked Variants Only” and “Non-Masked Variants Only” |
|  | VAF Minimum / Maximum | 0.01 / 1 |
| **Variant Annotation Filters** | SVs in ≤ this percentage of the control db samples with same enzyme | 100% |
|  | SV self-molecule check | SV found in self-molecules |
|  | Self-molecule count | Effective coverage x 0.025 |
|  | SVs in ≤ this percentage of the control db samples | 0%  100% (Dual Parental/iPSC Analysis) |
|  | SV chimeric score filter | Show not failing chimeric score |
|  | SV overlapping genes filter | All SVs |
|  | SV control sample assembly check | Not found in paired control assemblies |
|  | SV control sample molecule check | Not found in paired control molecules |

**Table S6: Comparing PBMC-iPSCs Clone to Clone Distribution of SVs Across 39 PBMC-iPSC Clones**

| **PBMC** | **Insertions** | **Deletions** | **Duplications** | **Translocations** |
| --- | --- | --- | --- | --- |
| SUSL 51 C4 | 0 | 0 | 0 | 0 |
| SUSL 62 C2 | 0 | 0 | 0 | 0 |
| SUSL 62 C3 | 0 | 0 | 0 | 0 |
| SUSL 64 C1 | 0 | 0 | 0 | 0 |
| SUSL 64 C2 | 0 | 0 | 0 | 0 |
| SUSL 70 C1 | 0 | 0 | 0 | 0 |
| SUSL 76 C1 | 0 | 0 | 0 | 0 |
| SUSL 77 C3 | 0 | 0 | 0 | 0 |
| SUSL 78 C11 | 0 | 0 | 0 | 0 |
| SUSL 78 C6 | 0 | 0 | 0 | 0 |
| SUSL 79 C1 | 0 | 0 | 0 | 0 |
| SUSL 79 C2 | 0 | 0 | 0 | 0 |
| SUSL 90 C3 | 0 | 0 | 0 | 0 |
| SUSL 79 C4 | 0 | 0 | 1 | 0 |
| SUSL 90 C6 | 0 | 1 | 0 | 0 |
| SUSL 90 C9 | 0 | 1 | 0 | 0 |
| SUSL 51 C1 | 0 | 1 | 0 | 0 |
| SUSL 63 C1 | 0 | 1 | 0 | 0 |
| SUSL 63 C2 | 0 | 1 | 0 | 0 |
| SUSL 71 C1 | 0 | 1 | 1 | 0 |
| SUSL 71 C2 | 0 | 1 | 0 | 0 |
| SUSL 71 C3 | 0 | 1 | 1 | 0 |
| SUSL 76 C3 | 0 | 1 | 0 | 0 |
| SUSL 50 C8 | 0 | 4 | 0 | 0 |
| SUSL 70 C3 | 0 | 4 | 0 | 0 |
| SUSL 51 C2 | 1 | 0 | 0 | 0 |
| SUSL 70 C2 | 1 | 0 | 0 | 0 |
| SUSL 77 C2 | 1 | 0 | 2 | 0 |
| SUSL 77 C4 | 1 | 0 | 0 | 0 |
| SUSL 88 C5 | 1 | 0 | 0 | 0 |
| SUSL 62 C1 | 1 | 2 | 0 | 0 |
| SUSL 78 C10 | 1 | 2 | 0 | 0 |
| SUSL 64 C4 | 1 | 5 | 0 | 2 |
| SUSL 76 C2 | 2 | 0 | 0 | 0 |
| SUSL 50 C7 | 2 | 1 | 0 | 0 |
| SUSL 63 C6 | 2 | 1 | 0 | 0 |
| SUSL 50 C1 | 4 | 0 | 0 | 0 |
| SUSL 88 C2 | 4 | 1 | 0 | 0 |
| SUSL 88 C7 | 6 | 5 | 0 | 0 |

**Table S7: Comparing FBC-iPSCs Clone to Clone Distribution of SVs Across 34 FBC-iPSC Clones**

| **FBC** | **Insertions** | **Deletions** | **Duplications** | **Translocations** |
| --- | --- | --- | --- | --- |
| SUSL 06 C1 | 0 | 0 | 0 | 0 |
| SUSL 12 B3 | 0 | 0 | 0 | 0 |
| SUSL 40 C1 | 0 | 0 | 0 | 0 |
| SUSL 40 C7 | 0 | 0 | 0 | 0 |
| SUSL 40 C8 | 0 | 0 | 0 | 0 |
| SUSL 13 C1 | 0 | 0 | 1 | 0 |
| SUSL 25 C2 | 0 | 0 | 1 | 0 |
| SUSL 12 C1 | 0 | 1 | 0 | 0 |
| SUSL 13 C3 | 0 | 1 | 0 | 0 |
| SUSL 38 C2 | 0 | 1 | 0 | 0 |
| SUSL 13 C2 | 0 | 1 | 1 | 0 |
| SUSL 66 C4 | 0 | 1 | 1 | 0 |
| SUSL 25 C6 | 0 | 1 | 2 | 0 |
| SUSL 39 C2 | 0 | 2 | 1 | 0 |
| SUSL 45 C6 | 0 | 2 | 2 | 0 |
| SUSL 66 C5 | 0 | 3 | 0 | 1 |
| SUSL 07 C3 | 0 | 3 | 1 | 0 |
| SUSL 39 C8 | 1 | 0 | 0 | 0 |
| SUSL 06 C2 | 1 | 1 | 0 | 0 |
| SUSL 07 C5 | 1 | 1 | 0 | 0 |
| SUSL 07 C9 | 1 | 1 | 0 | 0 |
| SUSL 66 C2 | 1 | 1 | 1 | 0 |
| SUSL 39 C4 | 1 | 1 | 2 | 0 |
| SUSL 46 C2 | 1 | 1 | 2 | 0 |
| SUSL 06 C3 | 1 | 2 | 0 | 0 |
| SUSL 38 C1 | 1 | 2 | 0 | 0 |
| SUSL 46 C1 | 1 | 2 | 2 | 2 |
| SUSL 38 C5 | 1 | 4 | 0 | 0 |
| SUSL 45 C3 | 2 | 1 | 1 | 0 |
| SUSL 46 C9 | 2 | 1 | 4 | 0 |
| SUSL 73 C8 | 2 | 2 | 3 | 0 |
| SUSL 12 A2 | 3 | 1 | 1 | 0 |
| SUSL 73 C10 | 3 | 1 | 7 | 3 |
| SUSL 73 C3 | 3 | 3 | 0 | 0 |

**Table S8: 38 Acrocentric P arms (13p, 14p, 15p, 21p, and 22p) Removed from Downstream Analysis**

| **SUSL Line** | **Cell Type** | **Chromosome** | **SV Start (Label)** | **SV End (Label)** | **SV Size (bp)** | **SV Type** | **VAF** | **% In Control Database** | **Parental Molecule Count** | **Self-Molecule Count** |
| --- | --- | --- | --- | --- | --- | --- | --- | --- | --- | --- |
| SUSL 78 C6 | PBMC | 13p11.2 | 11,737,297 | 11,758,829 | 57,849 | Insertion | 0.27 | 16.1 | 4 | 11 |
| SUSL 88 C2 | PBMC | 13p11.2 | 11,737,297 | 11,758,829 | 10,241 | Insertion | 0.39 | 13 | 0 | 35 |
| SUSL 88 C2 | PBMC | 13p11.2 | 15,145,466 | 15,256,098 | 14,121 | Deletion | 1 | 0 | 0 | 15 |
| SUSL 88 C7 | PBMC | 13p11.2 | 11,469,987 | 11,494,953 | 9,118 | Insertion | 0.03 | 30.2 | 4 | 13 |
| SUSL 88 C7 | PBMC | 13p11.2 | 15,145,466 | 15,256,098 | 16,600 | Deletion | 1 | 0 | 0 | 16 |
| SUSL 88 C7 | PBMC | 13p11.2 | 11,117,365 | 11,214,165 | 31,078 | Insertion | 0.17 | 29.5 | 0 | 14 |
| SUSL 88 C7 | PBMC | 13p11.2 | 11,737,297 | 11,758,829 | 59,496 | Insertion | 0.15 | 16.1 | 0 | 13 |
| SUSL 12 B3 | FBC | 13p11.2 | 10,623,913 | 10,763,256 | 103,510 | Insertion | 0.16 | 9.1 | 3 | 13 |
| SUSL 40 C1 | FBC | 13p11.2 | 10,151,401 | 10,152,476 | 12,961 | Insertion | 0.25 | 3.5 | 2 | 13 |
| SUSL 13 C2 | FBC | 13p12 | 5,932,211 | 5,947,940 | 5,524 | Deletion | 0.4 | 0.4 | 0 | 26 |
| SUSL 13 C2 | FBC | 13p12 | 6,073,121 | 6,088,850 | 5,568 | Deletion | 0.43 | 0 | 1 | 21 |
| SUSL 13 C2 | FBC | 13p12 | 5,979,181 | 6,041,880 | 14,927 | Deletion | 0.37 | 0.7 | 0 | 23 |
| SUSL 12 A2 | FBC | 14p11.2 | 6,173,761 | 6,255,347 | 19,325 | Insertion | 0.26 | 0.7 | 3 | 12 |
| SUSL 12 A2 | FBC | 14p11.2 | 6,294,220 | 6,443,498 | 128,350 | Insertion | 0.39 | 0.4 | 2 | 12 |
| SUSL 73 C10 | FBC | 14p11.2 | 4,327,136 | 4,387,862 | 8,391 | Insertion | 0.2 | 5.6 | 4 | 12 |
| SUSL 64 C2 | PBMC | 14p11.2 | 7,962,696 | 8,045,596 | 14,193 | Insertion | 0.26 | 36.1 | 4 | 12 |
| SUSL 77 C2 | PBMC | 14p11.2 | 8,729,344 | 8,772,345 | 2,156 | Deletion | 0.1 | 0 | 2 | 13 |
| SUSL 88 C7 | PBMC | 14p11.2 | 8,085,093 | 8,118,049 | 3,310 | Insertion | 0.23 | 0 | 0 | 18 |
| SUSL 88 C7 | PBMC | 14p11.2 | 8,836,238 | 8,899,388 | 11,597 | Deletion | 0.34 | 14.4 | 2 | 31 |
| SUSL 88 C7 | PBMC | 14p11.2 | 7,962,696 | 8,045,596 | 17,883 | Insertion | 0.26 | 36.8 | 0 | 18 |
| SUSL 63 C6 | PBMC | 15p11.2 | 15,182,781 | 15,269,153 | 11,454 | Deletion | 1 | 43.9 | 3 | 12 |
| SUSL 50 C1 | PBMC | 15p12 | 4,042,293 | 4,056,850 | 8,849 | Insertion | 0.03 | 2.5 | 3 | 12 |
| SUSL 73 C3 | FBC | 15p12 | 4,532,091 | 4,536,512 | 25,251 | Insertion | 0.14 | 4.6 | 0 | 29 |
| SUSL 07 C3 | FBC | 15p13 | 57,089 | 71,189 | 2,938 | Insertion | 0.04 | 6.3 | 2 | 13 |
| SUSL 62 C1 | PBMC | 15p13 | 2,052,955 | 2,096,544 | 24,492 | Insertion | 0.1 | 3.2 | 0 | 64 |
| SUSL 78 C6 | PBMC | 21p11.2 | 5636126.5 | 5646566 | 3,181 | Deletion | 0.01 | 3.5 | 4 | 33 |
| SUSL 88 C5 | PBMC | 21p11.2 | 9,975,996 | 10,014,983 | 8,910 | Deletion | 0.71 | 29.5 | 4 | 46 |
| SUSL 88 C7 | PBMC | 21p11.2 | 8,433,366 | 8,449,992 | 3,115 | Insertion | 0.1 | 2.5 | 1 | 23 |
| SUSL 78 C11 | PBMC | 21p13 | 54003 | 68118 | 31,798 | Insertion | 0.01 | 0.7 | 3 | 16 |
| SUSL 50 C1 | PBMC | 22p11.2 | 10,297,596 | 10,451,423 | 131,265 | Deletion | 0.28 | 9.5 | 1 | 19 |
| SUSL 12 B3 | FBC | 21p11.2 | 6,027,695 | 7,794,379 | 457,026 | Deletion | 0.26 | 1.4 | 3 | 16 |
| SUSL 07 C9 | FBC | 21p12 | 3,168,573 | 3,172,998 | 3,131 | Insertion | 0.29 | 2.8 | 0 | 33 |
| SUSL 63 C6 | PBMC | 22p11.2 | 11,704,972 | 11,957,158 | 70,043 | Insertion | 0.29 | 1.1 | 3 | 13 |
| SUSL 88 C5 | PBMC | 22p11.2 | 10,711,477 | 11,009,102 | 48,710 | Deletion | 0.17 | 12.3 | 0 | 14 |
| SUSL 88 C5 | PBMC | 22p11.2 | 12,412,572 | 12,442,697 | 73,478 | Insertion | 0.27 | 14.7 | 2 | 16 |
| SUSL 88 C7 | PBMC | 22p11.2 | 5,934,964 | 6,062,323 | 39,536 | Insertion | 0.2 | 11.2 | 0 | 17 |
| SUSL 78 C6 | PBMC | 22p13 | 158994 | 197921 | 3,408 | Insertion | 0.02 | 6.7 | 0 | 14 |
| SUSL 06 C1 | FBC | 22p13 | 55,682 | 69,787 | 3,382 | Deletion | 0.01 | 18.2 | 3 | 13 |
